# Evolution of the canonical sex chromosomes of the guppy and its relatives

**DOI:** 10.1101/2020.09.25.314112

**Authors:** M. Kirkpatrick, J.M. Sardell, B. J. Pinto, G. Dixon, C. L. Peichel, M. Schartl

## Abstract

The sex chromosomes of the guppy, *Poecilia reticulata*, and its close relatives are of particular interest: they are much younger than the highly degenerate sex chromosomes of model systems such as mammals and *Drosophila melanogaster*, and they carry many of the genes responsible for the males’ dramatic coloration. Over the last decade, several studies have analyzed these sex chromosomes using a variety of approaches including sequencing genomes and transcriptomes, cytology, and linkage mapping. Conflicting conclusions have emerged, in particular concerning the history of the sex chromosomes and the evolution of suppressed recombination between the X and Y. Here we address these controversies by reviewing the evidence and reanalyzing data. We find no support for a nonrecombining sex determining region (SDR) or evolutionary strata in *P. reticulata*. Further, we find that the evidence most strongly support the hypothesis that the SDRs of *P. picta* and *P. wingei* evolved independently after those lineages diverged. We identify possible causes of conflicting results in previous studies and suggest best practices going forward.

## 1: INTRODUCTION

The origin and evolution of young sex chromosomes are of particular interest to evolutionary genomics. They are the most rapidly evolving part of the genome in many animals and plants, and they have features that give unique insights into the evolution of recombination, sexually antagonistic selection, and other important processes (Bachtrog *et al*. 2011). The guppy, *Poecilia reticulata*, holds a special place in the history of this subject. Because they carry many of the genes responsible for the males’ famed coloration, the *P. reticulata* sex chromosomes have been studied since the 1920s (Winge 1922; Haskins *et al*. 1961; Lindholm and Breden 2002; Charlesworth 2018). The last decade has seen a burst of research on the sex chromosomes of the guppy and its relatives. Several recent studies have arrived at conflicting conclusions, notably regarding the evolution of recombination. These controversies are the focus of this paper.

Studies from the pre-genomic era also provided conflicting conclusions regarding recombination between the sex chromosomes. Using linkage maps, Tripathi *et al*. (2009) reported that recombination between the X and Y of *P. reticulata* is confined to a relatively small region bounded on either side by large nonrecombining regions. Nanda *et al*. (2014) used cytology and linkage maps to study the sex chromosomes of *P. reticulata*, its sister species *P. wingei*, and the closely related *P. obscura*. They concluded that the Y chromosomes of these three species descended from a common ancestral Y, and that the X and Y chromosomes of *P. reticulata* recombine down their lengths (save perhaps for a small heterochromatic region specific to the Y). Further, they reported that an extended region of heterochromatin at the end of the Y distal to the centromere has evolved in *P. wingei* that would regionally block recombination with the X. Conversely, by inferring meiotic crossovers visualized through the localization of MHL1, Lisachov *et al*. (2015) found that recombination between the X and Y in *P. reticulata* is concentrated towards the end of the chromosome distal to the centromere.

The pace of discovery accelerated with the arrival of genome sequences for *P. reticulata* (Fraser *et al*. 2015; Künstner *et al*. 2016). By analyzing molecular variation among re-sequenced genomes from several natural populations of *P. reticulata*, Wright *et al*. (2017) drew conclusions at odds with the previous reports. They reported that crossing over between the X and Y is completely blocked over about 40% (10 Mb) of their lengths, with recombining regions on either side. They claimed that the nonrecombining sex determining region, or SDR, is divided into two “strata”, which are regions in which crossing over between the X and Y was completely suppressed at different times in the past (Lahn and Page 1999; Charlesworth 2017). Wright *et al*. (2017) further concluded that the nonrecombining SDR has expanded independently in three populations that inhabit the headwaters of rainforest streams. Morris *et al*. (2018) followed up this study by identifying 40 loci that are unique to the putative nonrecombining region of the Y chromosome, and proposed two of them as candidates for the sex determining gene.

In the next study from the same research team, Darolti *et al*. (2019) enlarged the phylogenetic picture by analyzing genomes and transcriptomes from five species of poeciliid fish, including *P. reticulata*, its sister species *P. wingei*, and their congener *P. picta*. These authors found that Chromosome 12 is responsible for sex determination in all three species. They reported that the two strata that they found on the *P. reticulata* Y are shared with *P. wingei*, which implies that they evolved in the common ancestor of those species. In the more distantly related *P. picta*, they found the Y chromosome to be highly degenerate. They also concluded that the SDRs in all three species descend from a common ancestor, which implies that the rates at which the Y degenerates varies greatly between lineages. Darolti *et al*. (2020) found from linkage mapping that completely suppressed recombination is confined only to the first of the two strata in *P. reticulata*, while there is very rare crossing over between the X and Y in the second “stratum”. In the most recent paper from that research group, Almeida *et al*. (2021) carried out long-read sequencing on much larger samples of individuals from six natural populations. They again concluded that the l data show that there is no recombination in Stratum 1.

An independent research team studied the *P. reticulata* sex chromosomes using linkage mapping (Bergero *et al*. 2019). They found no evidence for completely suppressed recombination between the X and Y anywhere on those chromosomes. Recombination between the X and Y in natural populations is very rare, however, as there is elevated *F*_ST_ between males and females down the entire length of Chromosome 12, in some regions attaining values greater than 0.1. Bergero *et al*. (2019) showed that the very low levels of recombination between the X and Y are not unique to the sex chromosomes: males also have extremely low recombination rates on autosomes except near the telomeres. They questioned the conclusion of Wright *et al*. (2017) that recombination between the X and Y has been suppressed. Bergero *et al*. also identified a region of high recombination on the end of the Y chromosome distal to the centromere, consistent with the patterns of male recombination on autosomes and the findings of Lisachov *et al*. (2015). The localization of crossovers in males to the tips of all chromosomes is consistent with the high GC content found there (Charlesworth *et al*. 2020b). Charlesworth *et al*. (2020a) reported a crossover between the X and Y of *P. reticulata* very close to the boundaries between Strata 1 and 2 as defined Almeida *et al*. (2021). Most recently, Fraser *et al*. (2020) assembled a new high quality reference genome for *P. reticulata* that includes unphased scaffolds from both the X and Y, and they re-sequenced fish from six natural populations. They concurred with Bergero *et al*. (2019) that there is no evidence for a nonrecombining stratum several megabases in size. Further, they found two candidate regions for the sex determining gene contained in Y-specific regions that (surprisingly) are located at opposite ends of the Y chromosome. Figure 1 summarizes some of these results.

**Figure 1:**
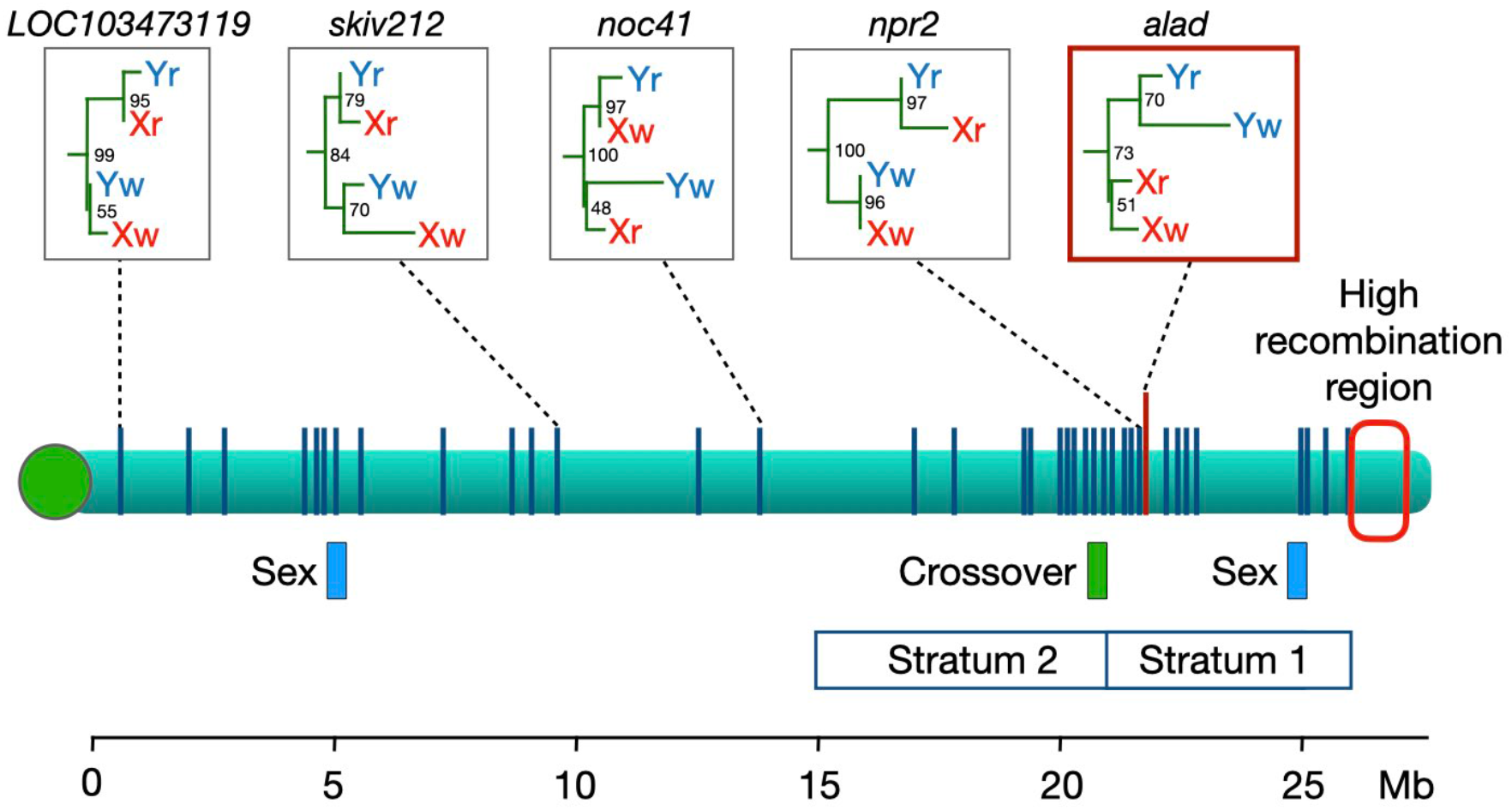
The sex chromosome of the guppy, *P. reticulata*. The horizontal green bar shows the interval in which a crossover was observed by Bergero *et al*. (2019; D. Charlesworth pers. comm.). Stratum 1 and Stratum 2 are regions where Wright *et al*. (2017) reported that the X and Y do not recombine. Blue boxes labeled “Sex” are candidate regions for the male-determining factor (Fraser *et al*. 2020). At far right is a region with a high local recombination rate (Lisachov *et al*. 2015, Bergero *et al*. 2019, Darolti *et al*. 2019). Vertical blue lines show locations of the 42 loci at which gene trees from *P. reticulata* (Xr, Yr) and *P. wingei* (Xw, Yw) were estimated by Darolti *et al*. (2020). Only the gene tree highlighted in the red box has a topology consistent with an ancestral SDR shared by the two species; examples of four other representative gene trees are also shown. Numbers at their nodes give the bootstrap support. The centromere is shown as the green circle at left.

These conflicting conclusions have led to controversy and confusion in the scientific community (Bergero and Charlesworth 2019; Wright *et al*. 2019). In an effort to remedy this situation, in this paper we review, reanalyze, and reinterpret published data from the guppy and its close relatives. Our goal is not to perform a forensic analysis to determine why different research teams have come to different conclusions. Rather, we analyze the existing data in order to independently evaluate conclusions regarding the evolutionary histories of their sex chromosomes.

We first focus on the evolution of recombination and the origin of the Y chromosomes. We find no evidence for strata in *P. reticulata* or for any region where crossing over between the X and Y is completely suppressed. Our results confirm that a non-recombining SDR spans much of the X and Y in *P. wingei* and *P. picta*, and that the Y in *P. picta* is highly degenerate.

We then consider hypotheses regarding homology of the SDRs in all three species. By “homology”, we mean that the SDRs in these species descended from a common ancestral SDR.

While the shared identity of the Y chromosomes of *P. reticulata* and *P. wingei* has been established by molecular cytogenetics (Nanda *et al*. 2014), that approach cannot evaluate the homology of their SDRs. To date, no genetic evidence has been adduced regarding homology of the Y chromosome or the SDR in *P. picta*. Our analysis of gene trees does not provide support for the hypothesis that the SDRs of *P. wingei* and *P. picta* descend from a common ancestral SDR. A second hypothesis is that the Y chromosomes of *P. wingei* and *P. reticulata* originated in their common ancestor when a new Y chromosome evolved from an X chromosome after divergence from *P. picta* (Bergero and Charlesworth 2019; Meisel 2020). This would make their Y chromosomes younger, and so explain why they are so much less degenerate than the Y of *P. picta*. But again, we find that gene trees give little support to this hypothesis. The most strongly supported hypothesis is that the SDRs of *P. wingei* and *P. picta* originated independently. We are unable to determine whether the entire Y chromosome originated once in the common ancestor of all three species and degenerated at different rates, as suggested by parsimony, or if the variation in Y degeneration across species reflects different ages of Y chromosomes that originated independently.

## 2: MATERIALS AND METHODS

### 2.1: The data

We downloaded all of the whole genome re-sequencing data available as of June 2020 from the NCBI SRA under Bioprojects PRJNA528814 and PRJNA353986. These data were acquired by Wright *et al*. (2017) and Darolti *et al*. (2019). (We did not use the more recent data from Fraser *et al*. (2020)) or Almeida *et al*. (2021) for most of our analyses because they were published after our paper was first submitted.) The data consist of paired-end Illumina sequences from 52 samples of whole genome sequences of 5 species: *Gambusia holbrooki, Poecilia latipinna, P. picta, P. reticulata*, and *P. wingei*. For a given data type, multiple records from the same individual were concatenated. Read quality was assessed using FastQC [v0.11.5] (Andrews 2010).

For most analyses, we mapped WGS reads to the *Xiphophorus maculatus* reference genome (v5.0; GCA_002775205.2; Schartl *et al*. (2013)) because it is the highest-quality poeciliid fish genome and is an equal phylogenetic distance from the *Poecilia* species. For the analyses of gene trees (see below) we redid our original analyses using the new *P. reticulata* reference genome (GCA_904066995.1) which was published after our paper was first submitted. We used this second genome with the goal of increasing read mapping rates across *Poecilia* species. Indeed, we found that the mapping efficiency with sequencing reads from *P. reticulata* is more than twice as high when using the new *P. reticulata* reference (∼93%) as when using the *X. maculatus* reference (∼41%), and it was similar across all three focal taxa.

However, there are no differences in read mapping efficiency between males and females using either of the two reference genomes, which makes *X. maculatus* an appropriate reference for the other analyses. There are small differences in the analysis pipelines using the *X. maculatus* and *P. reticulata* reference genomes that are specified below.

There is some uncertainty about the physical coordinates on the *P. reticulata* sex chromosomes. Different studies have used different reference genomes that vary in important features (e.g. Wright *et al*. (2017) *vs*. Bergero *et al*. (2019) *vs*. Fraser *et al*. (2020) *vs*. this study). Further, there are several reports of inversions and assembly errors on the *P. reticulata* sex chromosomes (Nanda *et al*. 2014; Bergero *et al*. 2019; Charlesworth *et al*. 2020a; Darolti *et al*. 2020; Fraser *et al*. 2020). Accordingly, we interpret the chromosome coordinates with caution.

#### Analyses based on the *X. maculatus* genome

We trimmed adapters and low-quality bases from raw SRA sequence reads using cutadapt [v1.14] (Martin 2011), mapped DNA and RNA reads to the *X. maculatus* reference genome using Bowtie2 [v2.3.4] with default parameters and the –local argument (Langmead and Salzberg 2012). We removed PCR duplicates using Picard [v2.21] (Broad Institute 2019). We sorted and subsetted alignment files by chromosome using SAMtools [v1.6] (Li *et al*. 2009) and calculated fold coverage in 10 kb windows using BEDtools [v2.26.0] (Quinlan and Hall 2010). We called variants with a quality score ≥20 using BCFtools mpileup [v1.10] (Li 2011). We removed indels and singletons, selected for biallelic SNPs using VCFtools [v0.1.16] (Danecek *et al*. 2011), and we required a minimum read depth of 3.

To determine which SNPs may be in an SDR, we considered patterns of heterozygosity. An allele was considered to be putatively Y-linked if it always appeared in males in heterozygotes and was absent from females. We designate these alleles as “Y-like”, the alternate alleles at these SNPs as “X-like”, and the sites at which these occur as “SDR-like SNPs”. We imposed the additional criterion that these sites have data from at least two individuals of each sex. Because the sample sizes are small, SDR-like SNPs can occur by chance at autosomal loci. Nevertheless, this set of sites will be highly enriched for SNPs that truly are sex-linked. As an internal control, we also applied this analysis to the autosomes to assess how frequently SNPs with putative sex linkage occur throughout the genome.

To study divergence between the X and Y chromosomes, we computed five statistics in sliding windows: (1) the ratio of read depth in males and females (“read depth ratio”), computed using BEDtools [v2.26.0] (Quinlan and Hall 2010); (2) *F*_ST_ between males and females (“*F*_ST_”), based on Weir and Cockerham’s (1984) estimator computed using VCFtools [v0.1.16] (Danecek *et al*. 2011); (3) the ratio of the density of all SNPs in males and in females (“SNP density ratio”), computed using VCFtools and a custom R script; (4) the density of SNPs with patterns of heterozygosity consistent with the SDR (“SDR-like SNP density”), computed using VCFtools and a custom R script; and (5) the density of SNPs with alleles that are restricted to females (“female-specific SNP density”), computed using VCFtools and a custom R script. (The latter are unexpected in species with XY sex determination, but we will see below that they do appear in one of the species.) Because all downstream analyses are dependent on successfully mapping reads, we first calculated read depth differences between males and females in highly-sensitive 10 kb windows. All other sliding-window statistics we calculated in 100 kb windows.

#### Analyses based on the *P. reticulata* genome

We quality and adapter trimmed the sequencing reads using Trim Galore! [v0.6.6] (Andrews 2010; Martin 2011), filtered PCR duplicates using bbmap [v38.90] (Bushnell 2014), and mapped reads to the *P. reticulata* reference genome using minimap2 [2.17] (Li 2018). We called variants one chromosome at a time at sites with quality score ≥20 using BCFtools mpileup [v1.11] (L
I 2011). We removed indels and singletons, selected for biallelic SNPs using VCFtools [v0.1.16] (Danecek *et al*. 2011), and we required a minimum read depth of 10.

We implemented this strategy using a custom R script. We first split the genotypes by species using VCFtools. We then filtered SNPs to include only those with no missing individuals (--max-missing 1.0) and a minimum allele frequency of 0.2 (--maf 0.2). We then split the genotypes further by sex and calculated allele frequencies (--freq) and observed heterozygosity (--hardy) in VCFtools.

We used gene trees to test hypotheses regarding the evolution of suppressed recombination. We applied several filters with a custom R script to identify SNPs used to construct these trees. We refer to the SNPs that meet all of the following criteria as “topologically informative.” First, we required sequence from at least two *P. wingei* males, two *P. wingei* females, two *P. picta* males, two *P. picta* females, one *P. latpinna*, and one *G. holbrooki*. Second, we only considered SNPs where alleles are fixed within X and Y chromosomes. That is, we required that the females within a species are all homozygous for the same allele, and that the males in that species either are all heterozygous or all homozygous for the same allele found in females. Third, we required that all individuals from the outgroup species (*P. latpinna* and *G. holbrooki*) possess the same allele, which we identified as the “ancestral” allele. Finally, we required that the alternate “derived” allele is fixed in either two or three of the following four chromosomes: the *P. wingei* X, *P. wingei* Y, *P. picta* X, or *P. picta* Y. We used the same R script to calculate the number of topologically informative SNPs that exhibit each of the 10 gene trees shown in Figure 5.

## 3: RESULTS

### 3.1: Divergence and recombination between the X and Y

To quantify divergence between the X and Y in the three species, we examined several statistics in sliding windows using the syntenic linkage group 8 of *X. maculatus* as the reference. Figure 2 shows the read depth ratio and male:female *F*_ST_, while Figure 3 shows the male:female SNP density ratio, SDR-like SNP density, and female-specific SNP density. Figure 4 shows the densities of SDR-like SNPs on the sex chromosomes and autosomes. As anticipated, SDR-like SNPs do occur on autosomes, but are much less common (see Methods).

**Figure 2.**
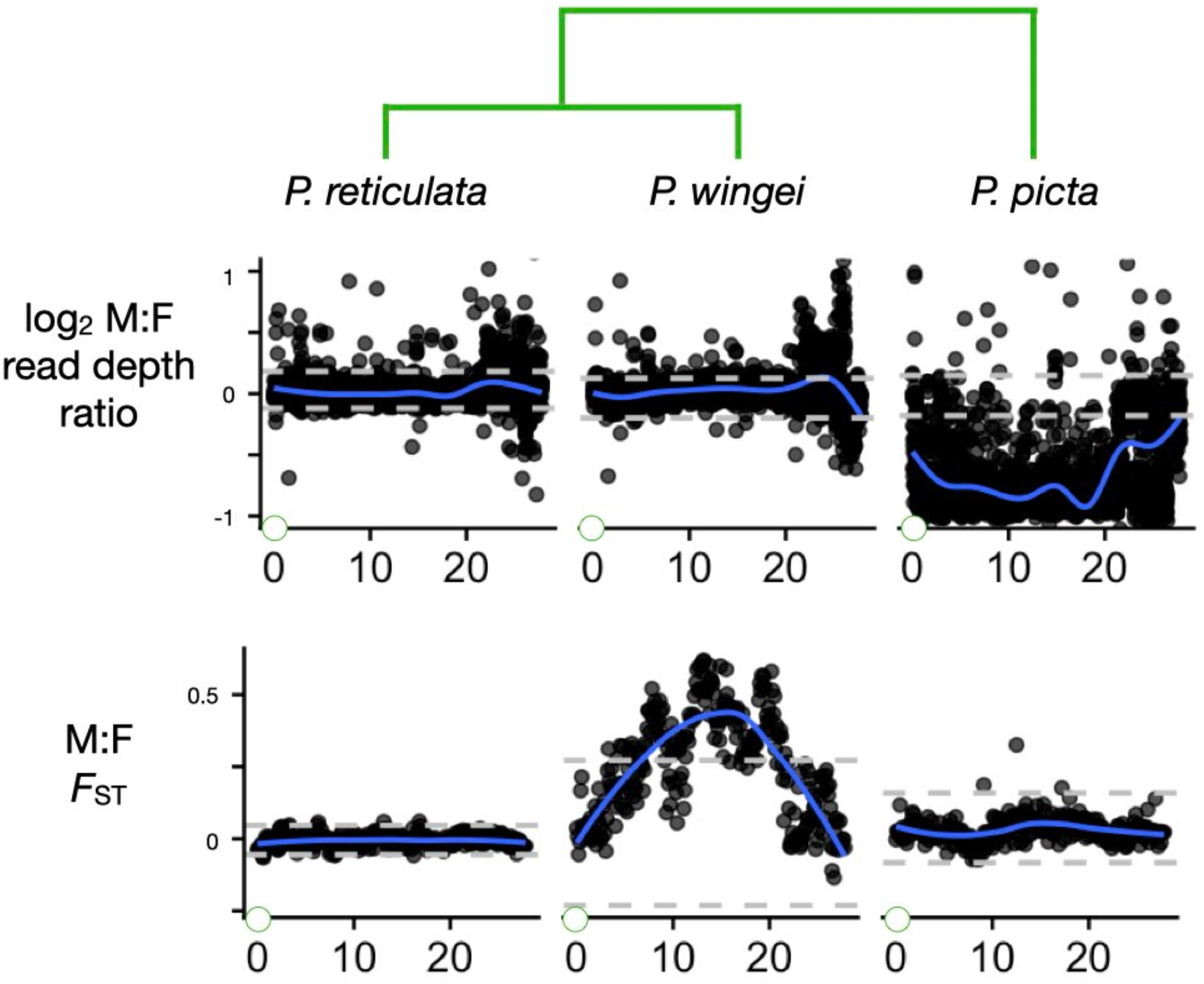
Divergence between the X and Y chromosomes in the three species as reflected by two statistics that measure differences between the sexes: the male:female read depth, and the male:female *F*_ST_. The gray horizontal dashed lines show the bottom 2.5% and top 97.5% intervals based on windows from all autosomes. The blue curves are smoothed regressions. Green circles at the left of the Y axes represent the centromere. The phylogeny is shown at top.

**Figure 3.**
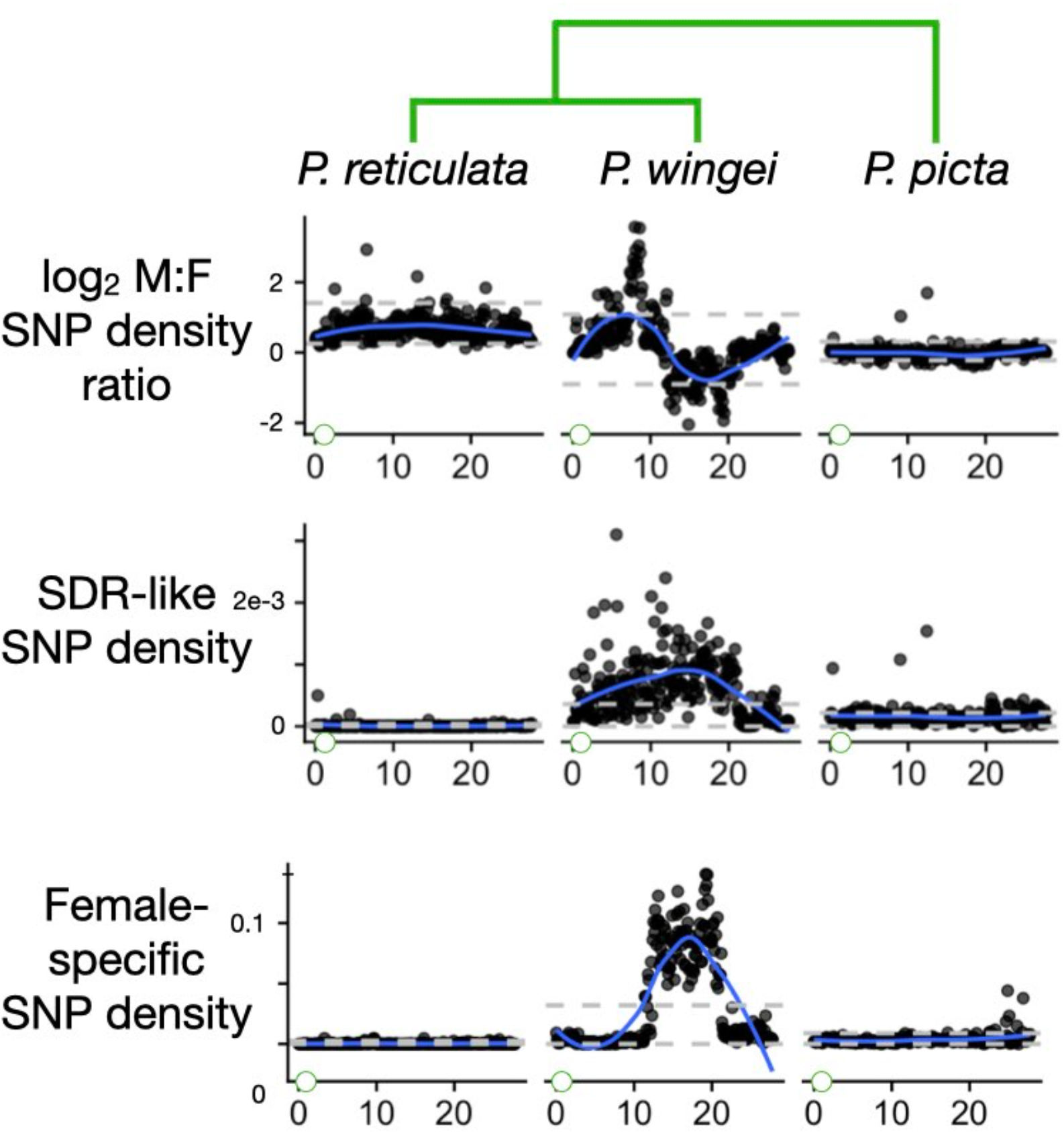
Divergence between the X and Y chromosomes as measured by three additional statistics: the male:female SNP density ratio, the SDR-like SNP density, and the female-specific SNP density. The gray horizontal dashed lines show the bottom 2.5% and top 97.5% intervals based on windows from all autosomes. The blue curves are smoothed regressions. Green circles at the left of the Y axes represent the centromere. The phylogeny is shown at top.

**Figure 4.**
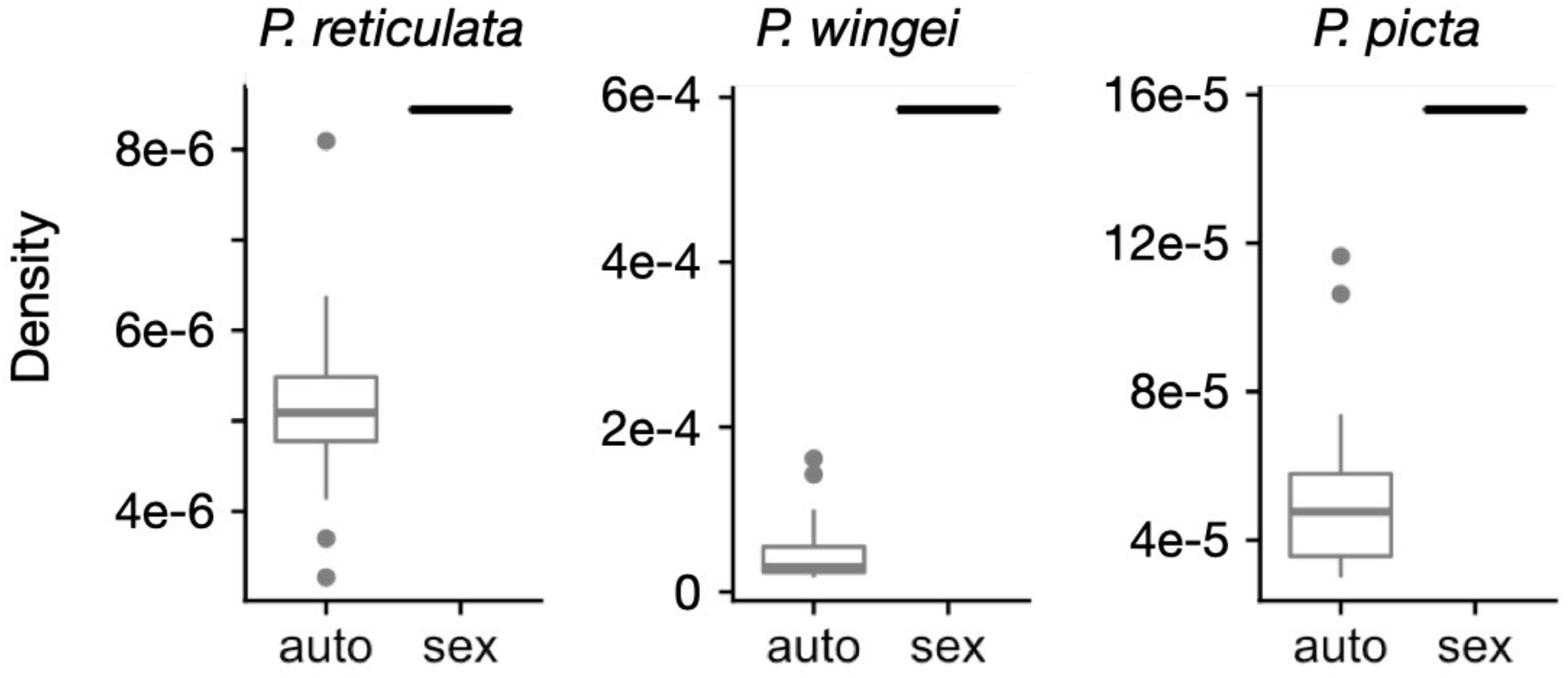
Densities per base pair of SDR-like SNPs (see Methods) that map to the sex chromosomes and to the autosomes.

**Figure 5.**
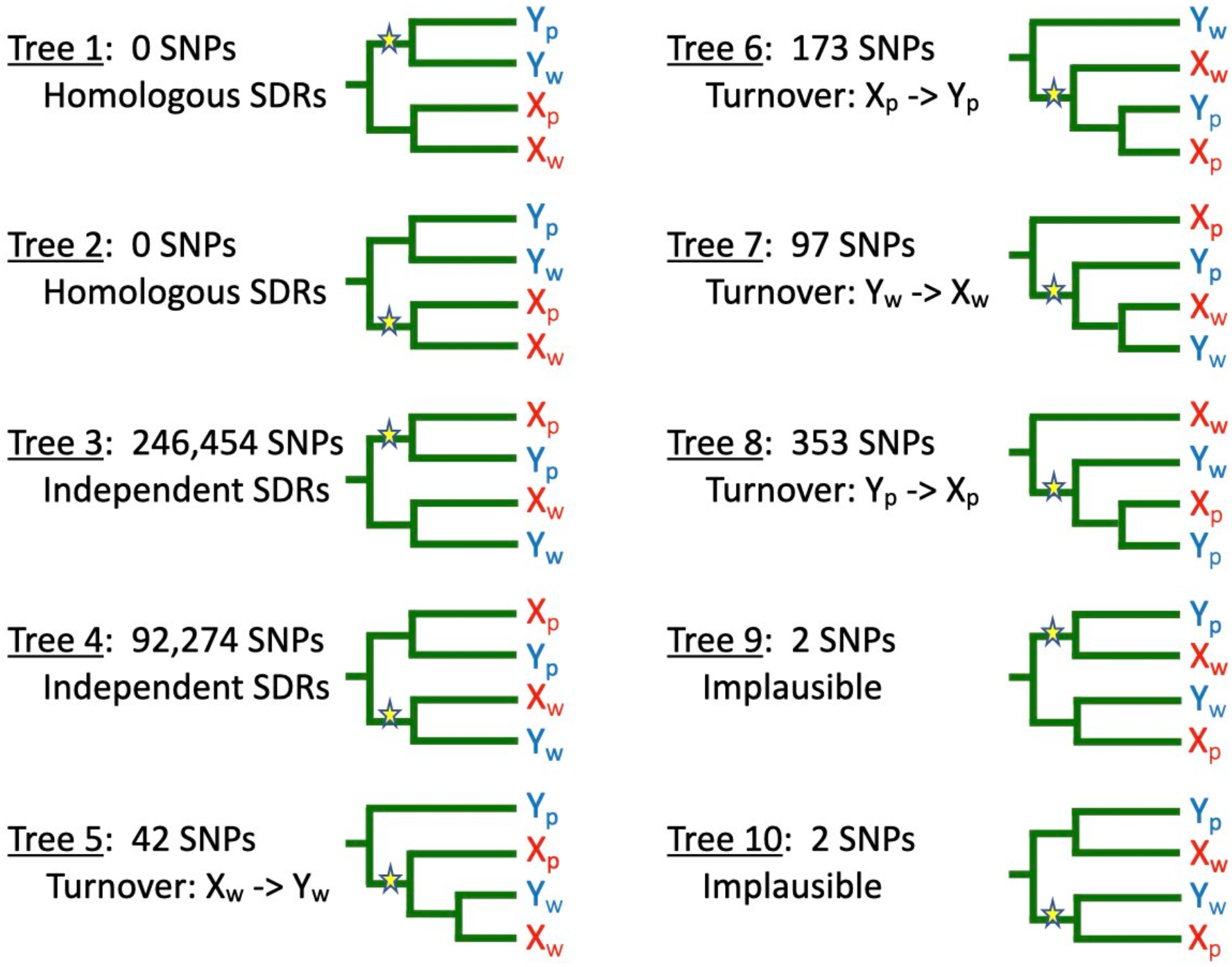
Ten trees for the SDRs of *P. picta* and *P. wingei* inferred from allele sharing patterns at topologically informative SNPs. X_p_ and Y_p_ represent the sex chromosomes of *P. picta*; X_w_ and Y_w_ represent the sex chromosomes of *P. wingei*. Derived mutations are shown by stars. Indicated are the numbers of SNPs at which each tree is seen and a biological interpretation of that tree. All trees can also result from introgression, homoplasy, and sequencing error. Note that the subtrees descending from the mutations shown in Trees 5 – 8 are consistent with the proposed biological scenario, but the allele sharing patterns observed at those SNPs give no information about the evolutionary relations of the three types of sex chromosomes that carry the mutant allele.

In *P. reticulata*, there is no sign that the X and Y have diverged very much anywhere along their lengths. We do not see a decreased read depth ratio, which is a telltale signature of Y chromosome degeneration (Vicoso and Bachtrog 2015). The maximum value of the smoothed regression for *F*_ST_ between males and females is less than 0.003 down the entire length of the sex chromosomes (Figure 2). The maximum value in any 100 kb window is *F*_ST_ = 0.15. (We obtained similar results for *F*_ST_ using both the *P. reticulata* and the *X. maculatus* reference genomes.) The other four statistics fall within the ranges typical for autosomes. These results do not provide any evidence of a nonrecombining sex determining region (SDR).

Our results are consistent with the results but not the interpretation of previous analyses based on gene trees (Darolti *et al*. 2020; Almeida *et al*. 2021). Following the origin of a non-recombining stratum on the Y by an inversion (or any other mechanism involving a *cis* recombination modifier), the SDRs on all Y chromosomes that inherit that stratum will form a monophyletic group with respect to the X chromosomes (Lahn and Page 1999; Zhou *et al*. 2014).

The monophyly persists through speciation events: the homologous strata on the Ys from all the descendant species continue to form a clade with respect to the X chromosomes, and the monophyly extends across the entire length of the stratum. In recombining regions of the sex chromosomes, however, this monophyly breaks down. Recombination causes gene copies from the X and Y chromosomes of each species to be intermingled on a gene tree. Recombination events more recent than a speciation event cause gene copies from the X and Y of a species to cluster together rather than with their gametologs in other species. Gene trees thus offer a sensitive way to distinguish the SDR from the recombining pseudoautosomal region (PAR) because they integrate signals of recombination that have accumulated in natural populations over many generations (Dixon *et al*. 2018; Sardell *et al*. 2021).

Darolti *et al*. (2020) estimated the gene trees at 42 loci spread along the length of the sex chromosomes. Figure 1 shows their locations. Only one of these loci, *alad*, shows a gene tree topology consistent with a nonrecombining stratum shared between *P. reticulata* and *P. wingei*, but the statistical support for this tree is weak (bootstrap value: 51%). (Three other gene trees are also stated to be consistent with that hypothesis, but in fact their topologies are not: see Figures 2c-e in Darolti *et al*. (2020).) At the locus *npr2*, which is less than 3 kb away from *alad*, there is strong statistical support for the node joining the X and Y in *P. reticulata* together (bootstrap value: 97%) and for a node joining the X and Y in *P. wingei* (bootstrap value: 96%).

This is evidence in both species that there has been recombination between the X and Y chromosomes in the region between the *npr2* locus and the sex determining region since the two species diverged. The remaining 40 gene trees are also inconsistent with a stratum that predates the divergence of *P. reticulata* and *P. wingei*. Most recently, Almeida *et al*. (2021) analyzed gene trees with much larger samples. In five of the six populations studied, the majority of trees in the proposed Stratum 1 and elsewhere on Chromosome 12 have topologies that are consistent with ongoing recombination between the X and Y in *P. reticulata*.

In the species that is sister to guppies, *P. wingei*, we find quite a different picture. Four of the five statistics (all but read depth ratio) fall far outside the autosomal range of values over much of the center part of the sex chromosomes (Figures 2 and 3). These patterns are consistent with very little or no recombination. The read depth ratio remains near 1, however, suggesting that there has not been extensive degeneration on the Y chromosome, possibly because recombination was suppressed recently. These conclusions concur with Darolti *et al*. (2019).

Within the region of reduced recombination in *P. wingei*, there is a smaller segment between about 10 Mb and 20 Mb where the female-specific allele density is greatly elevated, with very sharp boundaries at each end (Figure 3). As noted earlier, this pattern in unexpected in species with XY sex determination. The other three statistics just mentioned show no unusual patterns specific to this 10 Mb region. We speculate that these patterns might result from a polymorphic inversion on the X chromosome of *P. wingei* that is evident in the cytological data of Nanda *et al*. (2014).

The third species, *P. picta*, shows yet another distinctive set of patterns. Most strikingly, the read depth ratio is about one half over most of the end of the sex chromosome proximal to the centromere (Figure 2). This is indicative of large-scale deletions and/or divergence of the Y sequence to the point that reads from it no longer map to the reference (Vicoso 2019). The SDR-like SNP density, female-specific allele density, and *F*_ST_ are much lower than in *P. wingei*, presumably for the same reasons. These conclusions agree with Darolti *et al*. (2019).

### 3.2: Evolutionary relations between the SDRs

We used gene trees to investigate the evolutionary relations between the SDRs on the Y chromosomes of *P. wingei* and *P. picta*. One hypothesis is that the oldest parts of their SDRs are homologous, meaning that they descend from an SDR in their common ancestor. This possibility is favored by Darolti *et al*. (2019). A second hypothesis is that the Y chromosome of the shared ancestor of *P. wingei* and *P. reticulata* was derived from its X, an idea favored by Bergero and Charlesworth (2019) and Meisel (2020). A third hypothesis is that the SDRs of *P. wingei* and *P. picta* originated independently, either due to the independent evolution of suppressed recombination of a largely recombining ancestral Y, or independent recruitment of the same autosome as a sex chromosome.

These hypotheses make contrasting predictions regarding the gene trees and corresponding patterns of allele sharing for SNPs in the SDRs. If the SDRs of *P. wingei* and *P. picta* descend from a common ancestral SDR, then the Y chromosomes from both species will cluster together to the exclusion of the X chromosomes (Trees 1 and 2 in Figure 5). If the SDRs evolved independently in the two species, then the X and Y of each species will cluster together (Trees 3 and 4 in Figure 5). If the Y of *P. wingei* was derived from its X chromosome after that lineage diverged from *P. picta*, then the X and Y chromosomes of *P. wingei* will cluster together and form a clade that is sister to the *P. picta* X chromosomes (Tree 5 in Figure 5). Other evolutionary histories involving turnover events in which a Y was derived from an X or *vice versa* result in three additional topologies (Trees 6 - 8 in Figure 5).

None of the 339,397 SNPs have either of the topologies expected if the SDRs of *P. picta* and *P. wingei* are homologous (Trees 1 and 2). Instead, *P. picta* and *P. wingei* are fixed for different alleles at 99.8% of the SNPs (Trees 3 and 4), suggesting that their SDRs evolved independently. Unfortunately, bioinformatic artifacts resulting from Y degeneration can cause these gene trees to be erroneously inferred. At hemizygous sites (where there is a deletion on the Y), BCFtools and other widely-used SNP calling software impute a homozygote, which implies the presence of a Y-linked allele that is the same as the X-linked allele. This artifact results in gene trees in which the X and Y chromosomes from *P. picta* appear to cluster together to the exclusion of the X and Y chromosomes of *P. wingei*, which can falsely suggest that their SDRs evolved independently. Thus, the ubiquity of Trees 3 and 4 could result from the extreme degeneration of the Y in *P. picta* rather than from the true evolutionary history. It is possible to use criteria such as read depth to identify and remove hemizygous sites, but these filters are not always accurate, especially with the very small sample size used in this study.

Further analysis suggests, however, that hemizygosity is unlikely to be responsible for all the SNPs associated Trees 3 and 4. Tree 3, in which the derived allele is unique to the X and Y chromosomes of *P. picta*, is 2.7 times more common than Tree 4, in which the derived allele is unique to the X and Y chromosomes of *P. wingei*. The predominance of Tree 3 is likely to be an artifact of degeneration in the Y of *P. picta*, as that tree will be erroneously inferred if a mutation that is unique to the X chromosome in *P. picta* occurs in a region deleted from the Y chromosome. Likewise, Tree 4 can be erroneously inferred because of the same bioinformatic error if the true evolutionary history is Tree 7. That evolutionary scenario is unlikely, however, since the origin of a Y from an X to our knowledge has never been reported. Tree 4 can also result from a mutation that is unique to the X chromosome of *P. wingei* in a region that is deleted from its Y chromosome. We found little evidence, however, of extensive Y degeneration in *P. wingei* based on the read depth ratio (Figure 2). These findings suggest that the large number of SNPs exhibiting Tree 4 (about 10% of all SNPs in our dataset, including non-fixed sites) likely reflect the true evolutionary history of the *Poecilia* sex chromosomes. This conclusion implies that the SDRs of *P. wingei* and *P. picta* originated independently.

The trees associated with sex chromosome turnover hypotheses (Trees 5 - 8 in Figure 5) are all very rare, and likely result from homoplasy and/or degeneration of the Y in *P. picta*. We first consider hypotheses based on homoplasy. At 42 SNPs, the gene tree is consistent the origin of a new Y chromosome from the X in the ancestor of *P. wingei* (Tree 5). But at more than twice as many SNPs, the X and Y of *P. wingei* both share a derived allele with the Y of *P. picta* (Tree 7). As explained above, this tree is consistent with an evolutionary hypothesis that is implausible. A more likely explanation is that Trees 5 and 7 arose when the same mutation independently occurred on one of the *P. picta* sex chromosomes and on a recombining region in *P. wingei* that later became part of the SDR. The greater frequency of Tree 7 relative to Tree 5 could be due to higher mutation rates on the Y than on the X in *P. picta*, as observed in species with male-biased mutation (Wilson Sayres and Makova 2011). Male-biased mutation would also explain the greater frequency of Tree 8 relative to Tree 6.

Now consider how degeneration of the Y in *P. picta* might lead to the patterns seen in Trees 5 – 8. Tree 5 can only be inferred at those rare sites in *picta* that are not hemizygous, while Tree 6 can be erroneously inferred at hemizygous sites. This contrast could explain why Tree 6 is much more common. The same logic could explain why Tree 8 is much more common than Tree 7. Finally, Trees 5 and 6 could also result if the SDRs of the two species are homologous and a SNP that evolved in the ancestral X occurs in a region that was lost from one of the Y chromosomes. We believe this hypothesis is unlikely as that scenario is incompatible with Trees 7 and 8, which are more common. Additionally, the SNPs showing Trees 5 and 6 occur throughout the chromosome rather than being colocalized in a region, as we expect for an ancestral SDR.

In sum, there is no evidence that supports the homology of the SDRs in *P. wingei* and *P. picta*. The gene trees instead suggest that these SDRs originated independently. Several details of the data can be understood as resulting from homoplasy and bioinformatic artifacts caused by the extreme degeneration of the Y in *P. picta*.

## 4: DISCUSSION

### 4.1: Current state of recombination on the X and Y chromosomes in the three species

The results presented above and those from previous studies provide several insights into the state of the sex chromosomes in these three species. In the guppy *P. reticulata*, five lines of evidence argue that the sex chromosomes have not evolved suppressed recombination or evolutionary strata and in fact continue to recombine in the region near to the sex determining gene (see Figure 1).

First, we see no signs of increased divergence between males and females (a proxy for divergence between the X and Y) in the candidate regions for sex determination or anywhere else along the chromosome (Figures 2 and 3).

Second, gene trees on the sex chromosomes presented by Darolti *et al*. (2020) suggest that crossovers have occurred in the region identified as nonrecombining by Wright *et al*. (2017) and later studies by that research team (Morris *et al*. 2018; Darolti *et al*. 2019; Darolti *et al*. 2020; Almeida *et al*. 2021). We expect that the most recent recombinant Y chromosome was established within the last few thousand generations, based on the effective population size of Y chromosomes relative to autosomes and the small census population sizes of guppies (Fraser *et al*. 2015).

Third, a male crossover has been directly observed in a region that had previously been proposed as a nonrecombining stratum (Bergero *et al*. 2019; Charlesworth *et al*. 2020a) (see Figure 1). It occurred near the 21 Mb position, which is also the approximate boundary between Strata 1 and 2 as defined by Almeida *et al*. (2021). While that paper states there is no X-Y crossing over in Stratum 1, it does allow for the possibility of rare events in Stratum 2.

Fourth, SNPs with heterozygosity patterns consistent with sex linkage are scattered across the length of the sex chromosomes (Bergero *et al*. 2019; Darolti *et al*. 2020; Almeida *et al*. 2021), rather than being restricted to the proposed nonrecombining region.

Fifth and finally, Y haplotypes have high molecular diversity in guppies (Almeida *et al*. 2021). This is consistent with the ongoing introduction of genetic variation by recombination from the X to the Y. It does not seem, however, compatible with a hypothesis proposed by Almeida *et al*., which is that recombination in Stratum 1 was suppressed gradually. No matter what the mechanism (an inversion, change in chromatin structure, etc.), the fixation of any *cis* modifier that expands the SDR by so much as a single base pair eliminates all variation throughout the entire SDR of the chromosome on which that change occurred. If all the suppression results from inversion(s) on the X, then diversity would be reduced on that chromosome, but that is not what the data show (Fig. 3c in Almeida *et al*. (2021)). Variation could be maintained if the SDR expanded -- either gradually or rapidly -- by one more more unlinked (*trans*) recombination modifiers. We do not know, however, of any species in which that has been observed, nor of any genetic mechanism that has that effect.

Bergero *et al*. (2019) and Charlesworth *et al*. (2020a) found that all chromosomes in males have extremely low recombination rates over most of their length, consistent with earlier reports in guppies (Tripathi *et al*. 2009; Lisachov *et al*. 2015). They suggested that the very low rate of crossing over between the X and Y is a simple consequence of this situation. We concur. Apparently, recombination in males is sufficiently rare in at least some populations that the X and Y chromosomes have diverged at the molecular level, *e*.*g. F*_ST_ between males and females in the range 0.05 to 0.1 over large regions of the sex chromosomes (Bergero *et al*. 2019; Almeida *et al*. 2021). (For unknown reasons, our analyses show lower levels of differentiation between males and females, with no clear peaks that exceed values found on the autosomes (Figure 2).)

While the degree of sex bias in recombination (heterochiasmy) is extreme in guppies, qualitatively similar differences between males and females are seen across the eukaryotes (Sardell and Kirkpatrick 2020). In fact, there are examples of heterochiasmy even more extreme than guppies. Recombination along almost the entire lengths of all chromosomes is extremely rare in males of ranid frogs (Brelsford *et al*. 2016; Jeffries *et al*. 2018). As a result, *F*_ST_between males and females approaches its maximum possible value on the sex chromosomes in some populations (Toups *et al*. 2019).

Variation between populations in recombination rates on the sex chromosomes is a recurring theme in the guppy literature. Using laboratory crosses, Haskins *et al*. (1961) showed a significant difference between two populations in the recombination rate between the sex-linked locus *Sb* and the sex determining gene. Fraser *et al*. (2015) reported differences between populations in the linkage of certain color patterns to the X and Y chromosome, but their findings regard differences in the frequencies of color pattern alleles on the X and Y chromosomes rather than differences in recombination rates. In contrast, Wright *et al*. (2017) and Almeida *et al*. (2021) argue that differences between populations in patterns of molecular variation reflect differences in the actual rates of recombination between the X and Y.

We suggest that the very subtle differences they reported between populations in the degree of molecular differentiation between the guppy X and Y could occur even in the absence of differences in the recombination landscape. Consider the consequences of a rare crossover between the X and Y, for example near to the male determining gene. If the new Y haplotype spreads to high frequency by selection or drift, differentiation between the X and Y will be erased from the crossover breakpoint down the rest of the chromosome distal to the male determining gene. In the headwaters of the streams where they live, population sizes of *P. reticulata* are only a couple thousand individuals (Fraser *et al*. 2015). Consequently, crossovers between the X and Y may occur very infrequently, giving the X and Y time to diverge slightly before the next recombinant Y chromosome is established. Downstream populations are many times larger (Fraser *et al*. 2015). The result will be that recombinant Y chromosomes appear much more frequently in downstream populations and so divergence between the X and Y has less opportunity to build up.

A second factor that could contribute to patterns of molecular variation is sexually antagonistic selection, or SAS. More than 50 traits that are under sexual selection in male *P. reticulata* have been mapped to the sex chromosomes (Lindholm and Breden 2002). These are expected to generate peaks in the divergence between the X and Y in neutral genetic variation and can generate patterns that give the appearance of suppressed recombination (Charlesworth 2018; Bergero *et al*. 2019). Recombination in *P. reticulata* males is so rare that the map length of almost the entire Y chromosome is much less than 1 cM (Bergero *et al*. 2019). The small census population sizes suggest that many targets of SAS may lie within less than one ρ (= *N*_e_ *r*/2) of the sex determining factor, a condition favorable to inflated values of *F*_ST_between the X and Y (Kirkpatrick and Guerrero 2014). Indeed, peaks in *F*_ST_between males and females (a proxy for divergence between the X and Y) consistent with SAS have been reported by Bergero *et al*. (2019) and are visible in Figure 1 of Almeida *et al*. (2021). Thus differences between populations in the intensity of sexual selection and the frequencies of alleles at loci that experience SAS could also contribute to the differences between populations in the degree of X-Y divergence even in the absence of variation in recombination rates.

The recent publication of a greatly improved reference genome for *P. reticulata* that includes the Y (Fraser *et al*. 2020) adds additional insight. Those authors concluded that if an SDR exists, it is very small (less than 1 Mb). Of the two candidate regions for sex determination that they found, sequence from the most likely one is absent from the X chromosome. This would explain why it was not detected in studies (including ours) that are based on earlier reference genomes that lack Y-specific sequences. Analysis of gene trees might determine which of the two candidate regions identified by Fraser *et al*. (2020) harbors the sex determination gene.

In the guppy’s sister species, *P. wingei*, molecular divergence between males and females suggests that the X and Y have extremely low or no recombination over most of their lengths. Divergence is clearly evident in four of the five statistics shown in Figures 2 and 3.

Within that large region, there is a smaller segment of about 10 Mb that shows greatly elevated female-specific allele density. This region could reflect a polymorphic inversion on the X chromosome of *P. wingei* (Nanda *et al*. 2014). If so, the patterns seen in the figures could result if the three X chromosomes in our sample of *P. wingei* males are monomorphic for one arrangement while the six X chromosomes in the females include both arrangements. We are not able to determine if there is more than one stratum blocking recombination between the X and Y in *P. wingei*. A segment of the sex chromosomes distal to the centromere shows no sign of molecular divergence between the X and Y, suggesting high rates of recombination there.

The most dramatic patterns are seen in *P. picta*. Consistent with the findings of Darolti *et al*. (2019), our analyses suggest that most of its Y chromosome no longer recombines, and much of it has degenerated by sequence divergence or deletion (Figures 2 and 3).

### 4.2: History of the sex chromosomes and their SDRs

Our analysis of gene trees lead us to reject the hypotheses that the SDRs of *P. picta* and *P. wingei* are largely homologous. Although this conclusion is complicated by degeneration of the Y in *P. picta*, a study of *Gasterosteus* sticklebacks showed a clear signal of SDR homology using gene trees, even though the Y chromosome of one of the species shows about the same degree of degeneration as the Y of *P. picta* (Sardell *et al*. 2021). We likewise reject an alternative hypothesis is that a new Y was derived from its X in *P. wingei* after that lineage diverged from *P. picta*.

The most parsimonious explanation for the data is one originally proposed by Darolti *et al*. (2019): the guppy and its two congeners share an ancestral Y chromosome, one with an extremely small SDR (as in *P. reticulata*). This would explain the absence of signal of homology in our gene trees, as they can only detect homology of non-recombining regions of the Y chromosome. But in contrast to the conclusions of Darolti *et al*., we find that suppressed recombination between the X and Y most likely evolved independently in the ancestors of *P. picta* and *P. wingei*. Under this hypothesis, the Y may be much more degenerate in *P. picta* either because recombination between its X and Y was suppressed much earlier or because degeneration proceeded much more rapidly than in *P. wingei*. Both situations are seen in *Gasterosteus* sticklebacks (Sardell *et al*. 2021). The homology of the SDRs in *P. picta* and *P. wingei* might best be investigated by long-read sequencing technology to determine if these species share Y-specific sequences identified in the guppy by Fraser *et al*. (2020).

### 4.3: Reconciliation of past studies and best practices going forward

How can the many discrepancies between the conclusions from previous studies be reconciled, and how best can the field move forward? Some of the problems can be traced to semantic differences. We defer to the terminology for sex chromosomes defined by Bachtrog *et al*. (2011). The sex determining region, or SDR, is a segment of the sex chromosomes that includes the sex determining factor and in which crossing over between the X and Y is completely suppressed. Within the SDR, there can be one or more strata, which are regions in which crossing over was suppressed at different times in the past (Lahn and Page 1999; Charlesworth 2018). The SDR may be as small as a nucleotide, as in the case of the fugu, or as large as most of the Y chromosome, as in mammals (Bachtrog *et al*. 2014). By this definition, it is a *non sequitur* to say there is a stratum on the guppy sex chromosomes in which there is rare crossover recombination (Darolti *et al*. 2020; Almeida *et al*. 2021). Again following Bachtrog *et al*. (2011), all of the sex chromosomes that fall outside of the SDR make up the pseudoautosomal region, or PAR, regardless of whether the local recombination rate (measured as cM/Mb) is very high, as in mammals, or very low, as in some frogs (Bachtrog *et al*. 2014). By this definition, it is also a *non sequitur* to refer to only part of the recombining region as the PAR (Bergero *et al*. 2019). Authors are of course free to adopt noncanonical definitions, but in that case, we urge them to give explicit definitions and use terms consistently.

There are, however, three substantive reasons why different studies have arrived at differing conclusions. First, some statistical approaches used to define the SDRs and PARs of the guppy and its relatives are problematic. Two nonrecombining strata were identified in *P. reticulata* and *P. wingei* using the criterion that molecular differences between males and females over a region of the sex chromosome fall outside the 95% confidence interval seen on autosomes (Wright *et al*. 2017; Darolti *et al*. 2019). By that standard, 5% of the entire genome is expected to be identified as nonrecombining sex chromosome strata, and indeed several regions of autosomes do meet that criterion (Supplementary Figure 1 in Wright *et al*. 2017).

Further, the differences between males and females used to define nonrecombining strata in those studies are extremely small and likely not biologically meaningful (*e*.*g*. a male:female SNP density ratio differing much less than 1% from the autosomal average (Wright *et al*. 2017)).

Additionally, three studies have given unambiguous evidence that the Y chromosome of *P. reticulata* recombines in or very near the region that had been proposed as a nonrecombining stratum (Figure 1) (Bergero and Charlesworth 2019; Charlesworth *et al*. 2020a; Darolti *et al*. 2020). Thus the bioinformatic strategy used by Wright *et al*. (2017), Darolti *et al*. (2019), and Almeida *et al*. (2021) appears to be inadequate to identify SDRs and strata. Finally, although the authors state otherwise, the gene trees presented by Darolti *et. al*. (2020) provide strong evidence of ongoing recombination, as only one poorly-supported gene tree shows the topology expected for a non-recombining region.

Second, the results from linkage mapping experiments have been interpreted in different ways. It is difficult to draw conclusions from a failure to observe crossovers, especially in crosses with few offspring and when recombination is rare. Mapping experiments are underpowered to distinguish between partial and full linkage of loci to the male determining gene (Muyle *et al*. 2016; Wright *et al*. 2019). This limitation is particularly acute in species with extremely low recombination rates in males, such as the guppy. Conclusions about regions where the X and Y do not recombine based on linkage mapping should be treated with caution.

Methods that define nonrecombining sex determining regions using gene trees are much more sensitive because they integrate genetic signatures of recombination over long periods of evolutionary time (*e*.*g*. (Lahn and Page 1999; Zhou *et al*. 2014; Dixon *et al*. 2018; Sardell *et al*. 2021).

Third, sequencing studies have used different reference genomes and different strategies to map reads. Wright *et al*. (2017) used a *de novo* assembly of *P. reticulata* (N50 = 0.017 Mb) whose scaffolds were then oriented according to a guppy reference. Bergero *et al*. (2019) and Charlesworth *et al*. (2020a) mapped their DNA reads to the old *P. reticulata* reference (scaffold N50 = 31.4 Mb; Künstner *et al*. (2016)). While this reference has the advantage of being from the species of interest, it is based entirely on scaffolded short-read sequences. Darolti *et al*. (2019; 2020) used one of the publicly-available *Xiphophorus helleri* genomes (scaffold N50 = 29.4 Mb; Shen *et al*. (2016)) to order scaffolds from their own *de novo* genome assemblies of *P. reticulata*. Last, for most of the analyses in this paper we used the *Xiphophorus maculatus* reference because it has the highest quality of any species in this family (Schartl *et al*. 2013). The current assembly (version 5) is based on long-read as well as short-read sequencing and optical mapping to produce the chromosome-level assembly (scaffold N50 = 31.5 Mb). To verify our findings in *X. maculatus*, we re-analyzed the gene trees on Chr 12 using the new *P. reticulata* reference genome (scaffold N50 = 32.8 Mb) that was recently released by Fraser *et al*. (2020).

Many of the obstacles that have made studies of the guppy sex chromosomes so difficult will hopefully soon be in the past as the use of experimental phasing of sex chromosomes (Sardell *et al*. 2021) and advances in long-read sequencing eliminate many of the bioinformatic problems.

## ACKNOWLEDGEMENTS

We are grateful to members of the Kirkpatrick lab and two anonymous reviewers for discussion and comments. This study was supported by NIH grant R01-GM116853 to M.K..

## DATA ACCESSIBILITY

All the data used in the study were acquired from the publicly-accessible sources cited in the text. The custom scripts used for data processing, analysis, and figure generation are available on GitHub at https://github.com/grovesdixon/guppy_sex_chroms.

## AUTHOR CONTRIBUTIONS

M.K. conceived of the study. J.M.S., B.J.P., and G.D.D. conducted the analyses. All authors contributed to the writing.

## REFERENCES

Almeida, P., B. A. Sandkam, J. Morris, I. Darolti, F. Breden et al., 2021 Divergence and remarkable diversity of the Y chromosome in guppies. Molecular Biology and Evolution 38: 619–633.

Andrews, S., 2010 FastQC: a quality control tool for high throughput sequence data, pp.

Bachtrog, D., M. Kirkpatrick, J. E. Mank, S. F. McDaniel, J. C. Pires et al., 2011 Are all sex chromosomes created equal? Trends in Genetics 27: 350–357.

Bachtrog, D., J. Mank, C. Peichel, S. Otto, M. Kirkpatrick et al., 2014 Sex determination: Why so many ways of doing it? PLoS Biology 12: e1001899.

Bergero, R., and D. Charlesworth, 2019 Reply to Wright et al.: How to explain the absence of extensive Y-specific regions in the guppy sex chromosomes. Proceedings of the National Academy of Sciences of the United States of America 116: 12609–12610.

Bergero, R., J. Gardner, B. Bader, L. Yong and D. Charlesworth, 2019 Exaggerated heterochiasmy in a fish with sex-linked male coloration polymorphisms. Proceedings of the National Academy of Sciences of the United States of America 116: 6924–6931.

Brelsford, A., N. Rodrigues and N. Perrin, 2016 High-density linkage maps fail to detect any genetic component to sex determination in a Rana temporaria family. Journal of Evolutionary Biology 29: 220–225.

Broad Institute, 2019 Picard Toolkit.

Bushnell, B., 2014 BBMap: a fast, accurate, splice-aware aligner pp. Lawrence Berkeley National Laboratory, Berkeley, CA.

Charlesworth, D., 2017 Evolution of recombination rates between sex chromosomes. Philosophical Transactions of the Royal Society B-Biological Sciences 372: 20160456.

Charlesworth, D., 2018 The guppy sex chromosome system and the sexually antagonistic polymorphism hypothesis for Y chromosome recombination suppression. Genes 9: 264.

Charlesworth, D., R. Bergero, C. Graham, J. Gardner and L. Yong, 2020a Locating the sex determining region of linkage group 12 of guppy (Poecilia reticulata). G3-Genes Genomes Genetics 10: 3639–3649.

Charlesworth, D., Y. X. Zhang, R. Bergero, C. Graham, J. Gardner et al., 2020b Using GC content to compare recombination patterns on the sex chromosomes and autosomes of the guppy, Poecilia reticulata, and its close outgroup species. Molecular Biology and Evolution 37: 3550–3562.

Danecek, P., A. Auton, G. Abecasis, C. A. Albers, E. Banks et al., 2011 The variant call format and VCFtools. Bioinformatics 27: 2156–2158.

Darolti, I., A. E. Wright and J. E. Mank, 2020 Guppy Y chromosome integrity maintained by incomplete recombination suppression. Genome Biology and Evolution 12: 965–977.

Darolti, I., A. E. Wright, B. A. Sandkam, J. Morris, N. I. Bloch et al., 2019 Extreme heterogeneity in sex chromosome differentiation and dosage compensation in livebearers. Proceedings of the National Academy of Sciences of the United States of America 116: 19031–19036.

Dixon, G., J. Kitano and M. Kirkpatrick, 2018 The origin of a new sex chromosome by introgression between two stickleback fishes. Molecular Biology and Evolution 36: 28–38.

Fraser, B. A., A. Kunstner, D. N. Reznick, C. Dreyer and D. Weigel, 2015 Population genomics of natural and experimental populations of guppies (Poecilia reticulata). Molecular Ecology 24: 389–408.

Fraser, B. A., J. R. Whiting, J. R. Paris, C. J. Weadick, P. J. Parsons et al., 2020 Improved reference genome uncovers novel sex-linked regions in the guppy (Poecilia reticulata). Genome Biology and Evolution 12: 1789–1805.

Haskins, C. P., E. F. Haskins, J. J. A. McLaughlin and R. E. Hewitt, 1961 Polymorphism and population structure in Lebistes reticulatus, a population study, 320–395 in Vertebrate Speciation, edited by W. F. Blair. University of Texas Press, Austin.

Jeffries, D. L., G. Lavanchy, R. Sermier, M. J. Sredl, I. Miura et al., 2018 A rapid rate of sex-chromosome turnover and non-random transitions in true frogs. Nature Communications 9: 4088.

Kirkpatrick, M., and R. F. Guerrero, 2014 Signatures of sex-antagonistic selection on recombining sex chromosomes. Genetics 197: 531–541.

Künstner, A., M. Hoffmann, B. A. Fraser, V. A. Kottler, E. Sharma et al., 2016 The genome of the Trinidadian guppy, Poecilia reticulata, and variation in the Guanapo population. Plos One 11: e0169087.

Lahn, B. T., and D. C. Page, 1999 Four evolutionary strata on the human X chromosome (vol 286, pg 964, 1999). Science 286: 2273–2273.

Langmead, B., and S. L. Salzberg, 2012 Fast gapped-read alignment with Bowtie 2. Nature Methods 9: 357–359.

Li, H., 2011 A statistical framework for SNP calling, mutation discovery, association mapping and population genetical parameter estimation from sequencing data. Bioinformatics 27: 2987–2993.

Li, H., 2018 Minimap2: pairwise alignment for nucleotide sequences. Bioinformatics 34: 3094–3100.

Li, H., B. Handsaker, A. Wysoker, T. Fennell, J. Ruan et al., 2009 The sequence alignment/map format and SAMtools. Bioinformatics 25: 2078–2079.

Lindholm, A., and F. Breden, 2002 Sex chromosomes and sexual selection in poeciliid fishes. American Naturalist 160: S214–S224.

Lisachov, A. P., K. S. Zadesenets, N. B. Rubtsov and P. M. Borodin, 2015 Sex chromosome synapsis and recombination in male guppies. Zebrafish 12: 174–180.

Martin, M., 2011 Cutadapt removes adapter sequences. from high-throughput sequencing reads. EMBnet Journal 17: 10–12.

Meisel, R. P., 2020 Evolution of sex determination and sex chromosomes: a novel alternative paradigm. BioEssays 2020: 1900212.

Morris, J., I. Darolti, N. I. Bloch, A. E. Wright and J. E. Mank, 2018 Shared and species-specific patterns of nascent Y chromosome evolution in two guppy species. Genes 9: 238.

Muyle, A., J. Kafer, N. Zemp, S. Mousset, F. Picard et al., 2016 SEX-DETector: a probabilistic approach to study sex chromosomes in non-model organisms. Genome Biology and Evolution 8: 2530–2543.

Nanda, I., S. Schories, N. Tripathi, C. Dreyer, T. Haaf et al., 2014 Sex chromosome polymorphism in guppies. Chromosoma 123: 373–383.

Quinlan, A. R., and I. M. Hall, 2010 BEDTools: a flexible suite of utilities for comparing genomic features. Bioinformatics 26: 841–842.

Sardell, J. M., M. P. Josephson, A. C. Dalziel, C. L. Peichel and M. Kirkpatrick, 2021 Heterogeneous histories of recombination suppression on stickleback sex chromosomes. Molecular Biology and Evolution msab179.

Sardell, J. M., and M. Kirkpatrick, 2020 Sex differences in the recombination landscape. American Naturalist 195: 361–379.

Schartl, M., R. B. Walter, Y. J. Shen, T. Garcia, J. Catchen et al., 2013 The genome of the platyfish, Xiphophorus maculatus, provides insights into evolutionary adaptation and several complex traits. Nature Genetics 45: 567–572.

Shen, Y. J., D. Chalopin, T. Garcia, M. Boswell, W. Boswell et al., 2016 X. couchianus and X. hellerii genome models provide genomic variation insight among Xiphophorus species. BMC Genomics 17: 67.

Toups, M. A., N. Rodrigues, N. Perrin and M. Kirkpatrick, 2019 A reciprocal translocation radically reshapes sex-linked inheritance in the common frog. Molecular Ecology 28: 1877–1889.

Tripathi, N., M. Hoffmann, E.-M. Willing, C. Lanz, D. Weigel et al., 2009 Genetic linkage map of the guppy, Poecilia reticulata, and quantitative trait loci analysis of male size and colour variation. Proceedings of the Royal Society of London. Series B: Biological Sciences 276: 2195–2208.

Vicoso, B., 2019 Molecular and evolutionary dynamics of animal sex-chromosome turnover. Nature Ecology & Evolution 3: 1632–1641.

Vicoso, B., and D. Bachtrog, 2015 Numerous transitions of sex chromosomes in Diptera. Plos Biology 13: e1002078.

Weir, B. S., and C. C. Cockerham, 1984 Estimating F-statistics for the analysis of population structure. Evolution 38: 1358–1370.

Wilson Sayres, M. A., and K. D. Makova, 2011 Genome analyses substantiate male mutation bias in many species. BioEssays 33: 938–945.

Winge, O., 1922 One-sided masculine and sex-linked inheritance in Lebistes reticulatus. Journal of Genetics 12: 145–162.

Wright, A. E., I. Darolti, N. I. Bloch, V. Oostra, B. Sandkam et al., 2017 Convergent recombination suppression suggests role of sexual selection in guppy sex chromosome formation. Nature Communications 8: 14251.

Wright, A. E., I. Darolti, N. I. Bloch, V. Oostra, B. A. Sandkam et al., 2019 On the power to detect rare recombination events. Proceedings of the National Academy of Sciences of the United States of America 116: 12607–12608.

Zhou, Q., J. L. Zhang, D. Bachtrog, N. An, Q. F. Huang et al., 2014 Complex evolutionary trajectories of sex chromosomes across bird taxa. Science 346: 1246338.

